# Enabling CAR-T Cell Immunotherapy in Glioblastoma by Modifying Tumor Microenvironment via Oncolytic Adenovirus Encoding Bispecific T Cell Engager

**DOI:** 10.1101/2025.07.30.667708

**Authors:** Moon Jung Choi, Eui Young So, Bedia Akosman, Young Eun Lee, Alexander G. Raufi, Paul Bertone, Anthony M. Reginato, Clark C. Chen, Sean E. Lawler, Eric T. Wong, Olin D. Liang

## Abstract

Recent clinical trials show that CAR-T cell therapies can initially blunt tumor growth in glioblastoma (GBM) patients. However, the tumor microenvironment activates mechanisms that inhibit tumor-killing potential of the CAR-T cells and limit their therapeutic efficacy. To counteract this, we have utilized oncolytic adenovirus (OV) Ad5-Δ24-RGD as a platform to overexpress a bispecific T cell engager (BiTE) targeting both T cell marker CD3 and GBM specific tumor associated antigen IL-13Rα2. We first demonstrated that OV-BiTE could enhance recruitment of T cells to GBM *in vitro* and *in vivo*. We then showed that intratumoral injection of OV-BiTE followed by infusion of combined EGFR- and EGFRvIII-CAR-T cells was more effective than OV-BiTE supplemented with either CAR-T therapy alone, and led to significant tumor eradication in a GBM xenograft mouse model. In conclusion, our multimodal OV-BiTE & CAR-T cell immunotherapy is capable of overcoming immunosuppressive tumor microenvironment and GBM resistance to treatment.

**GRAPHICAL ABSTRACT:** 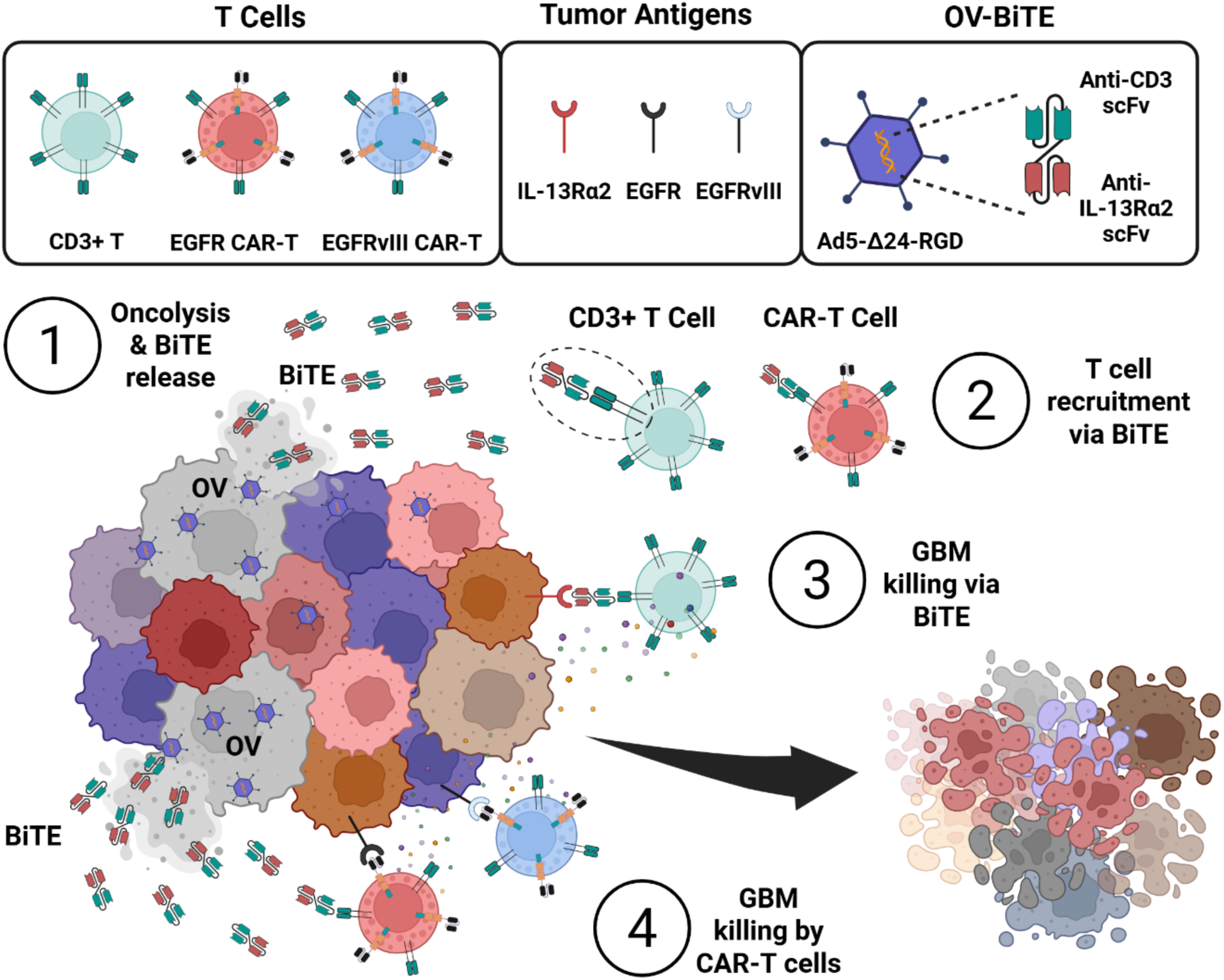

**HIGHLIGHTS:**

- Oncolytic adenovirus encoding bispecific T cell engager (OV-BiTE) combines two immunotherapeutic agents into one.
- OV-BiTE strategy modifies tumor microenvironment and enhances recruitment of T cells to glioblastoma (GBM) *in vitro* and *in vivo*.
- Multimodal OV-BiTE & CAR-T cell immunotherapy effectively reduced tumor mass in a GBM xenograft mouse model and is superior to either immunotherapy alone.

## Introduction

Glioblastoma (GBM) is the most lethal primary brain cancer, accounting for majority of all brain tumor cases ^1–4^. The average survival duration for GBM patients is 12-18 months after diagnosis, and the 5-year survival ratio is less than 5% of patients ^5,6^. Radiotherapy, chemotherapy, and surgical debulking remain the current standards of care for malignant glioma, but even tumor resection is difficult because of its location and potential neurological impairment ^7^. Its highly aggressive nature, molecular heterogeneity, the ability of resistant cancer stem cells to regrow post-therapy, the invasion of critical regions of the brain, and the inadequacy of achieving high therapeutic levels of chemotherapeutics in the brain because of the blood-brain barrier (BBB) are some of the key factors that constitute the vast amount of unmet need in GBM patients ^8^. Immunotherapeutic agents like immune checkpoint inhibitors (ICIs) ^9,10^, chimeric antigen receptor (CAR)-T cells ^11–13^, and oncolytic viruses ^14–16^ have revolutionized cancer therapy in some tumor types. However, their success in GBM remains limited because the GBM tumor microenvironment is highly immunosuppressive, rendering T cells incapable of recognizing and eliminating malignant cells ^17^. Hence, there is a critical need to develop effective immunotherapies for GBM.

CAR-T cell therapy uses autologous T cells engineered to target specific tumor antigens expressed on the surface of tumor cells for tumor eradication. CAR-T cell therapies have produced sustained therapeutic effects in refractory hematological cancers, but their success in the treatment of solid tumors has also been limited ^18–20^. The efficacy of CAR-T cells is restricted in GBM mainly because of the high degree of tumor heterogeneity, the BBB, and significantly immunosuppressive tumor microenvironment. Several CAR-T cell therapies targeting a range of tumor-associated antigens (TAAs) such as EGFR, EGFRvIII, IL13Rα2, HER2, B7-H3, and CD-147 are currently under clinical investigation in GBM ^21^. The loss of target antigen expression by glioma cells renders the CAR-T cell therapies ineffective, but targeting different tumor antigens can help overcome tumor resistance due to variable antigen expression ^20^. Three latest Phase I clinical trials have demonstrated feasibility, safety, and transient efficacy of intracerebroventricular delivered CAR-T cells for the treatment of GBM in patients ^22–24^. However, despite their clinical significance, there are limitations to the utility of CAR-T therapies ^25^. Collecting T cells from a patient is costly and time-consuming, especially for those with compromised immune systems. It is well-established that T cells are responsible for graft-versus-host disease in patients. Additionally, CAR-T cell therapy may cause severe side effects such as cytokine release syndrome or neurotoxicity. Finally, CAR-T cells struggle to penetrate the tumor microenvironment, which diminishes their therapeutic effectiveness. Hence, CAR-T cell therapy needs further optimization in GBM with improved accessibility to the brain, tumor cell targeting, survivability in immunosuppressive tumor microenvironment, and the ability to proliferate and exert therapeutic effects with minimal toxicities.

Oncolytic virus (OV) therapy uses replication-competent viruses that can selectively replicate and kill cancer cells. Direct oncolysis releases a wide range of TAAs, danger-associated molecular patterns and viral pathogen-associated molecular patterns, which trigger inflammatory immune responses in the tumor microenvironment ^26^. Most OV that are in clinical trials in GBM patients are being delivered locally to achieve an effective virus load in the tumors. A number of OV are being tested both at the preclinical and clinical level in GBM, among which some promising OV candidates are adenovirus [DNX-2401 (Ad5-Δ24-RGD)] ^14^, poliovirus (PVS-RIPO) ^27^, and retrovirus (Toca 511) ^28^. Ad5-Δ24-RGD is a conditionally replicative oncolytic adenovirus specifically engineered to destroy cancer cells. A 24-base pair deletion in the retinoblastoma (Rb)-binding domain of the E1A viral gene confers this adenovirus with selectivity to cancer cells ^29,30^, and an integrin binding RGD peptide has been incorporated into the H1 loop of the fiber to broaden tropism ^31^. Results from both preclinical ^32–37^ and clinical studies ^14,38–41^ indicate that Ad5-Δ24-RGD persists in human tumors for weeks, elicits tumor necrosis, triggers intratumoral immune cell infiltration, has no safety concerns, and can lead to long term patient benefit. However, following OV treatment, an antiviral immune response mediated by NK cells and macrophages may limit viral replication and mitigate the oncolytic effects. GBM has an “immune-cold” or noninflammatory tumor microenvironment ^42^ which has been challenging for monotherapies with ICIs or CAR T cells. Since OV can increase immune cell infiltration which could be crucial in breaking the immune tolerance and can improve tumor responsiveness to immunotherapies, Ad5-Δ24-RGD may serve as a primer for enhancing the efficacy of ICIs or CAR T cells. To this end, there is an ongoing clinical trial evaluating the combination of Ad5-Δ24-RGD and anti-PD1 antibody for GBM (NCT02798406). Also, combination of oncolytic adenovirus expressing IL-7 with B7H3-targeted CAR T cells showed higher efficacy than a single use of either of those treatments ^43^. In addition, the combination of OV with standard therapeutics like chemo- and radiotherapy can exert a synergistic effect in heterogeneous GBM ^44^. As a novel type of immunotherapy, bispecific T cell engager (BiTE) combines two single chain variable fragments (scFv) from different antibodies, one targeting CD3ε and the other targeting a TAA on the surface of tumor cells, which are connected by a short flexible linker. BiTE has unique therapeutic advantages as a specific tumor immunotherapy without the requirement of MHC-1 antigen presentation to activate T cells, form artificial immune synapses, and release perforin and granzyme to exert cytotoxic effects ^45^. When OV is used as a monotherapy, an antiviral immune response mediated by NK cells and macrophages may limit viral replication and mitigate the oncolytic effects. Thus, the synergy between OV and BiTE deserves special attention. OV can be used as a genetic engineering platform to express BiTE, which combines two different immunotherapies as one therapeutic agent. In addition, the selective replication of OV in tumors can limit the expression of BiTE in tumors, reducing the possibility of non-tumor side effects of BiTE.

The rationale for the current study is threefold: (i) To turn a non-responding “immune-cold” GBM tumor into a responsive “immune-hot” tumor through OV treatment; (ii) To recruit host cytotoxic T cells through BiTE to kill IL13Rα2-expressing GBM tumor cells in an MHC-1 independent manner; (iii) To enable infused CAR T cells to target GBM heterogeneity by killing EGFR- and EGFRviii-expressing tumor cells. Here, we show that intratumoral injection of OV-BiTE followed by infusion of combined EGFR- and EGFRvIII-CAR-T cells can lead to significant tumor eradication in a GBM xenograft mouse model, and this multipronged approach is superior to oncolytic or CAR-T immunotherapy alone, and. In conclusion, our multimodal OV-BiTE & CAR-T cell immunotherapy is capable of overcoming immunosuppressive tumor microenvironment and GBM resistance to treatment.

## Results

### Generation of CAR-T cells targeting GBM and profiling of CAR-T cell cytotoxic cytokines

We generated 3 CAR-T cells targeting GBM tumor antigens IL-13Rα2, EGFR and EGFRvIII, respectively. Briefly, three third-generation lentiviral CAR constructs ^46^ were made (Figure 1A and Figure S1): IL13Rα2-scFv-CD28-4-1BB-CD3ζ-EGFP, EGFR-scFv-CD28-4-1BB-CD3ζ-EGFP, and EGFRvIII-scFv-CD28-4-1BB-CD3ζ-EGFP. High titer lentiviruses were produced at Brown University Legorreta Cancer Center Cell & Genome Engineering Shared Resource, followed by transduction of primary CD8+ T cells isolated from human peripheral blood mononuclear cells. In addition to being frequently invoked previously by many investigators in their studies, these 3 GBM tumor antigens were also utilized in recent clinical trials ^22,24,47^, and were detectable via Western blots in commonly referenced GBM cell lines e.g., U87, U251 and U373, but not in normal human astrocytes (Figure 1B). Gene expression and surface expression of three target tumor antigens analyzed by PCR and flowcytometry (Figure S2). Fluorescence microscope and flow cytometry analyses indicated that T cell transduction efficiency in this study was approximately 35-40% (Figure 1C). Next, cytotoxic cytokine release profiles of CAR-T cells were evaluated in comparison with that of control CD8+ T cells using ELISPOT and ELISA assays, which were carried out following manufacturer’s instructions. For the ELISPOT assays, different groups of T cells were co-cultured with either U251 or U87 for 24 h, after which cytokine-positive spots for perforin, granzyme B and TNF-α were imaged and manually counted (Figure S3). When co-cultured with U87 GBM tumor cells, the production of both perforin and granzyme B was significantly higher in all 3 CAR-T groups than in the mock transduced T cell group, whereas proinflammatory cytokine TNF-α significantly increased only in the EGFR-CAR-T cell group (Figure 1D). When co-cultured with U251 GBM cells, the production of both TNF-α and granzyme B were significantly higher in CAR-T cell groups than in the mock transduced T cell group, whereas perforin significantly increased only in the IL13Rα2 group (Figure 1E). In summary, we have made 3^rd^ generation CAR-T cells which can recognize GBM specific tumor antigens and produce cytotoxic cytokines.

**Figure 1.**
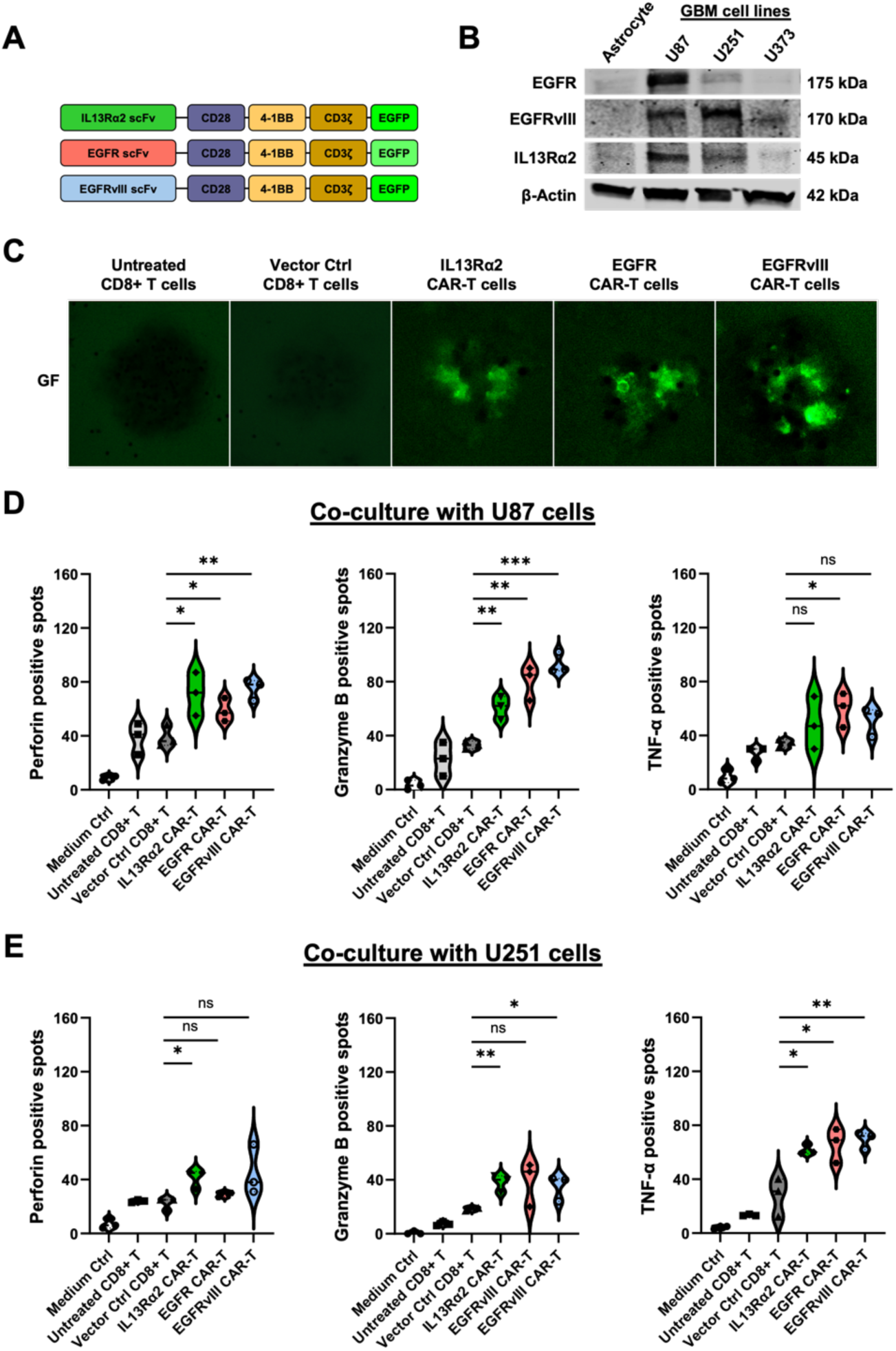
GBM-targeted CAR-T cells and cytokine release profiling. The representative CAR antigen (EGFR, EGFRviii, and IL13Rα2) expression in various GBM cell lines (U87, U251, and U373) GBM patient derived cell lines (GB2, GB11, and GB44), and hCMEC/D3 blood brain barrier endothelial cell line was analyzed by Flowcytometry (A). GFP on CAR-T cells detected by western blotting and microscopic analysis (B-C). And CAR-T cells were co-cultured with U87- and U251-MG cells for 24 hrs and their anti-cancer cytokines were detected by ELISPOT (D and E). Each value represents the mean ± S.D. for three separate experiments. *p < 0.05, ** p < 0.01, *** p < 0.001, and **** p<0.0001.

### *In vitro* 2D and 3D GBM tumor cell killing by CAR-T cells

Stable luciferase expressing U87 and U251 cell lines were generated through lentiviral transduction and puromycin selection of the GBM cells. For the 2D tumor cell killing assays, U87- and U251-luciferase expressing cells were co-cultured with medium alone, untreated CD8+ T cells, vector control CD8+ T cells, IL-13Rα2, EGFR or EGFRvIII CAR-T cells at an 1:1 effector-to-target (E:T) ratio (Figure 2A). The E:T co-culture conditions were experimentally determined separately (Figure S4). After 24 h of incubation, luciferase activities were quantified. As shown in Figure 2B, GBM tumor cell viability reduced significantly in all CAR-T cell treated groups in comparison with the control groups. In the 3D tumor cell killing assays ^48,49^, U87- and U251-spheroids were co-cultured with medium or various T cell groups as above, followed by propidium iodine (PI) staining for dead or dying cells. PI-positive cells were then visualized by using a fluorescence microscope and the mean fluorescence intensity (MFI) was quantified. In all CAR-T cell treated groups, the MFI values were significantly higher than those of the control groups (Figure 2C & D). Next, we also established a GBM blood-brain-barrier (BBB) spheroid model (Figure S5) to assess tumor killing ability of the CAR-T cells. As shown in Figure 2E, both EGFR and EGFRvIII CAR-T cells demonstrated significantly higher GBM tumor killing than CD8+ T cell controls. In summary, our IL-13Rα2, EGFR or EGFRvIII CAR-T cells can effectively kill GBM tumor cells in both 2D and 3D *in vitro* assays compared to CD8+ cytotoxicity T cells.

**Figure 2.**
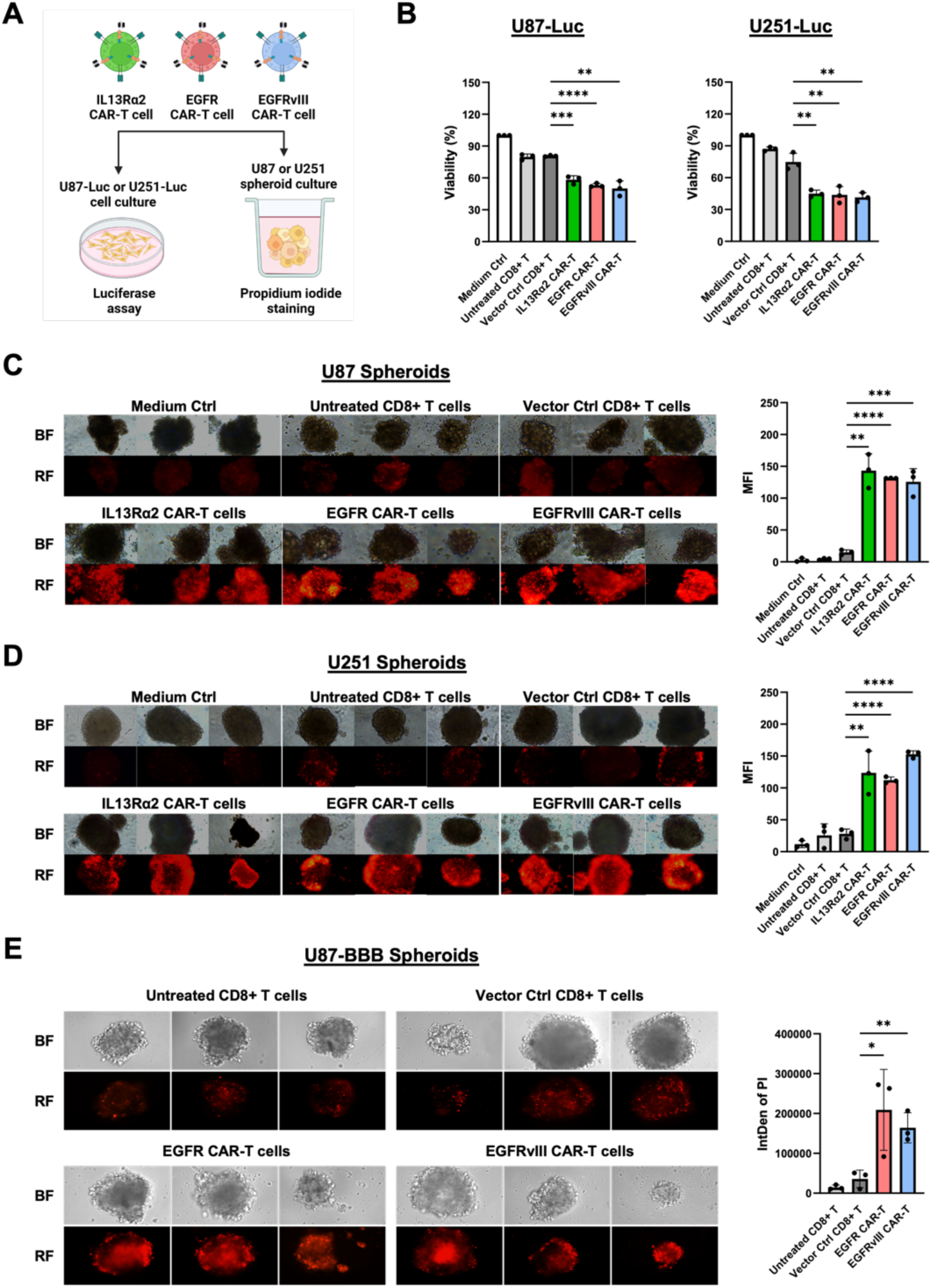
Anti-cancer killing of CAR-T cells in GBM cells and spheroids. Two GBM cells and spheroids were treated with three different CAR-T cells (A). The cell viability on the plate was analyzed by luciferase activity assay in U87-Luc and U251-Luc (B). After CAR-T cells treatment to GBM spheroids, the dead cells were detected with propidium iodine (PI) staining and analyzed by fluorescence microscope (10 X and 20 X) (C and D). And the anti-killing effect of CAR-T cells were analyzed in U87-BBB spheroids (E). Each value represents the mean ± S.D. for three separate experiments. ** p < 0.01, *** p < 0.001, and **** p<0.0001.

### Generation of oncolytic adenovirus Ad5-Δ24-RGD-BiTE, production of BiTE and oncolysis

We have adopted the AdenoQuick2.0 system (O.D.260 Inc, Boise, ID) to construct recombinant adenoviral vectors (Figure S6) and to produce Ad5-Δ24-RGD and Ad5-Δ24-RGD-BiTE adenoviruses. Figure 3A depicts the structure of BiTE, and its intended mode of action. The BiTE is a fusion protein comprising two single-chain variable fragments (scFv) of different antibodies on a single 55 kDa peptide ^50^, one of the scFv binds to T cells via CD3ε, and the other to a GBM specific tumor-associated antigen (TAA) IL13Rα2 ^12,51^. The two scFv were connected using a flexible 23 amino acids glycine/serine (G/S) linker in the following orientation: mAb OKT3 (VH)-mAb OKT3 (VL)-mAb IL13Rα2 (VL)-mAb IL13Rα2 (VH). Functionally, BiTEs have unique advantages as a specific tumor immunotherapeutic agent forming artificial immune synapses and promoting perforin and granzyme release by T cells to exert cytotoxic effects without the requirement of MHC-1 antigen presentation. In this study, we proposed that BiTE secretion from adenovirus infected GBM cells will build a concentration gradient which can recruit endogenous and therapeutic T cells into the tumor microenvironment. To demonstrate BiTE expression, adenoviral construct plasmid DNA was transfected into the 293FT cells and His-tagged BiTE protein was detected by Western blot using an antibody against the His tag (Figure 3B). To demonstrate oncolysis of GBM tumor cells by the adenoviruses, U87-Luc and U251-Luc cells and their respective spheroids were incubated with OV and OV-BiTE. Effective oncolysis of GBM tumor cells by the adenoviruses in 2D and 3D *in vitro* models was shown in Figure 3C & D. In summary, we have successfully generated oncolytic adenoviruses Ad5-Δ24-RGD and Ad5-Δ24-RGD-BiTE, where BiTE production by the latter construct was established. Furthermore, OV- and OV-BiTE-induced lysis of GBM tumor cells and spheroids *in vitro* was also demonstrated.

**Figure 3.**
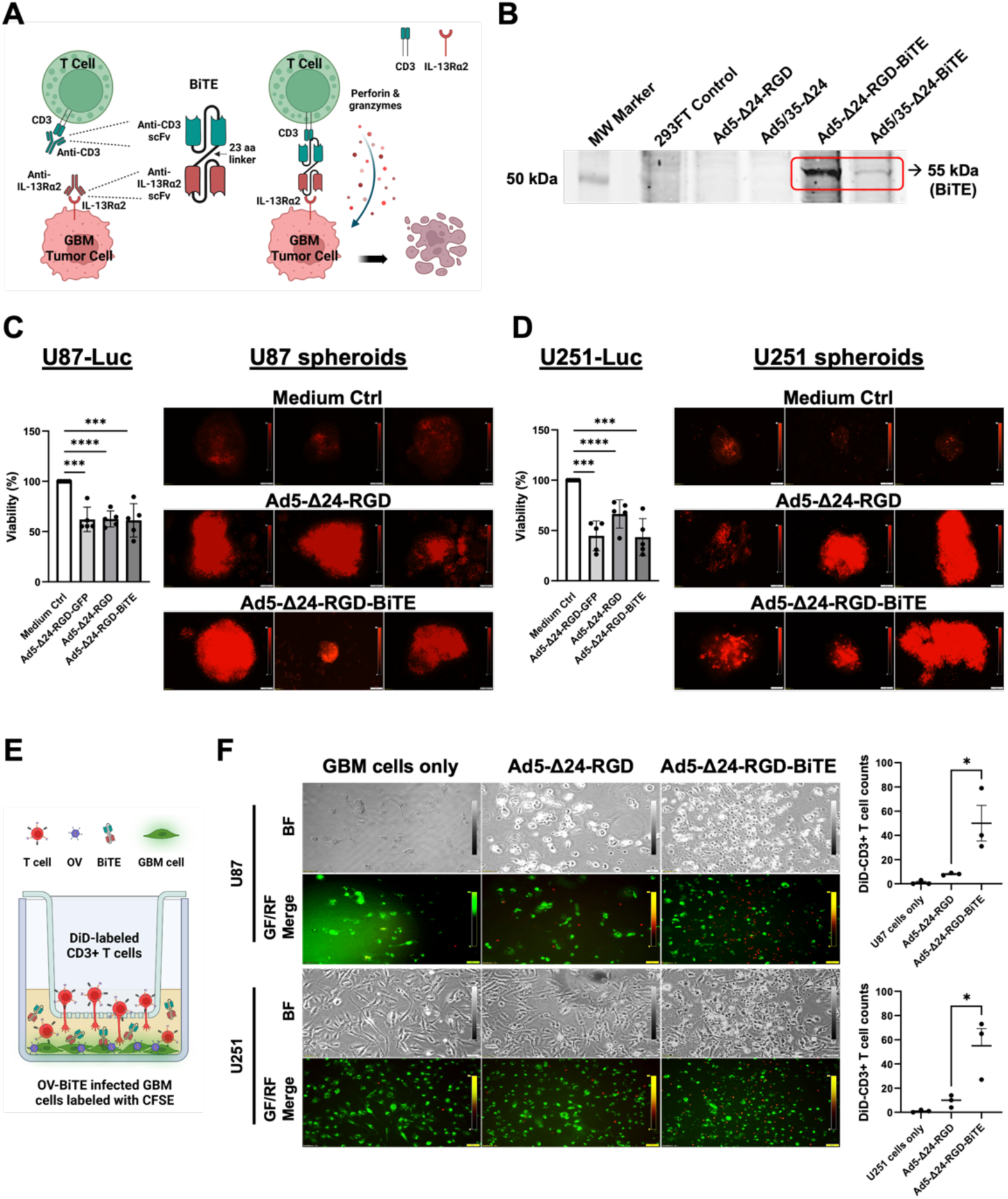
Anti-killing effect and in vitro T cell recruitment of oncolytic adenovirus (OAV) The BiTE protein that released from Ad-infected cancer cell can connect T cells and GBM cells (A). Adenoviral BiTE protein expression was determined by western blotting in 293FT cell (B). Oncolysis of OAV was analyzed by luciferase activity assay and PI staining in U87-luc and U251-luc (C and D). In vitro T cell recruitment assay of OAV was proceed transwell plate (E) and DiD-labeled T cells in lower chamber were detected with CSFE-labeled GBM cell lines by microscope analysis (F). Each value represents the mean ± S.D. for three separate experiments. *p < 0.05, *** p < 0.001, and **** p<0.0001.

### Enhanced T cell recruitment *in vitro* by OV-BiTE

Here, we conducted a transwell assay to demonstrate a enhanced recruitment of T cells *in vitro* by OV-BiTE compared to OV alone. As illustrated in Figure 3E, CFSE-labeled green GBM cells grown on the bottom of the lower chambers were infected by OV-BiTE or OV. Adenovirus containing cell culture medium was replaced with fresh medium after 4 h. DiD-labeled red CD3+ T cells were then added to the upper chamber of the transwell. After 24 h of incubation, the upper chambers were removed and the transwells were briefly centrifuged to pellet all the cells in the lower chamber. Both red and green fluorescence labeled cells on the bottom of the transwells were analyzed with a fluorescence microscope (Figure 3F). As a result, significantly more red fluorescent CD3+ T cells were detected in OV-BiTE infected group than in the OV infected group. In summary, we have demonstrated a significantly higher T cell recruitment *in vitro* by OV-BiTE treated GBM cells than OV alone treated or untreated GBM cells.

### Enhanced T cell recruitment *in vivo* by OV-BiTE

Next, we utilized a xenograft mouse model to determine whether the enhanced *in vitro* T cell recruitment by OV-BiTE over OV alone can be replicated *in vivo*. As illustrated in Figure 4A, GBM U87-Luc cells were subcutaneously injected into the right flank of immunodeficient mice, and the tumors were allowed to grow for 20 days followed by intratumoral injection of OV-BiTE or OV alone. Three days after adenoviral treatment, human CD3+ T cells were injected through the tail vein. Another 3 days later, tumor tissues were explanted for analysis. The apparent size and weight of isolated tumor masses from OV-BiTE or OV treated groups were significantly less than those of the untreated GBM controls, with OV-BiTE group exhibiting even greater tumor reduction than the OV group (Figure 4B & C). The tumor tissues were then homogenized into single cell suspensions and any infiltrated human CD3+ T cell populations were quantified per flow cytometry. Figure 4D & E show that OV-BiTE treated tumors had significantly higher numbers of human CD3+ T cells than those of the OV alone group. These findings were further supported by immunohistochemical staining of human CD3+ cells as dark brown spots in the tumor tissue sections. The numbers of human CD3+ cells in the OV-BiTE group were significantly higher than those from OV or untreated GBM control groups (Figure 4F & G). In summary, we have demonstrated a significantly higher T cell recruitment *in vivo* by OV-BiTE treated GBM tumors than OV alone treated or untreated GBM tumors.

**Figure 4.**
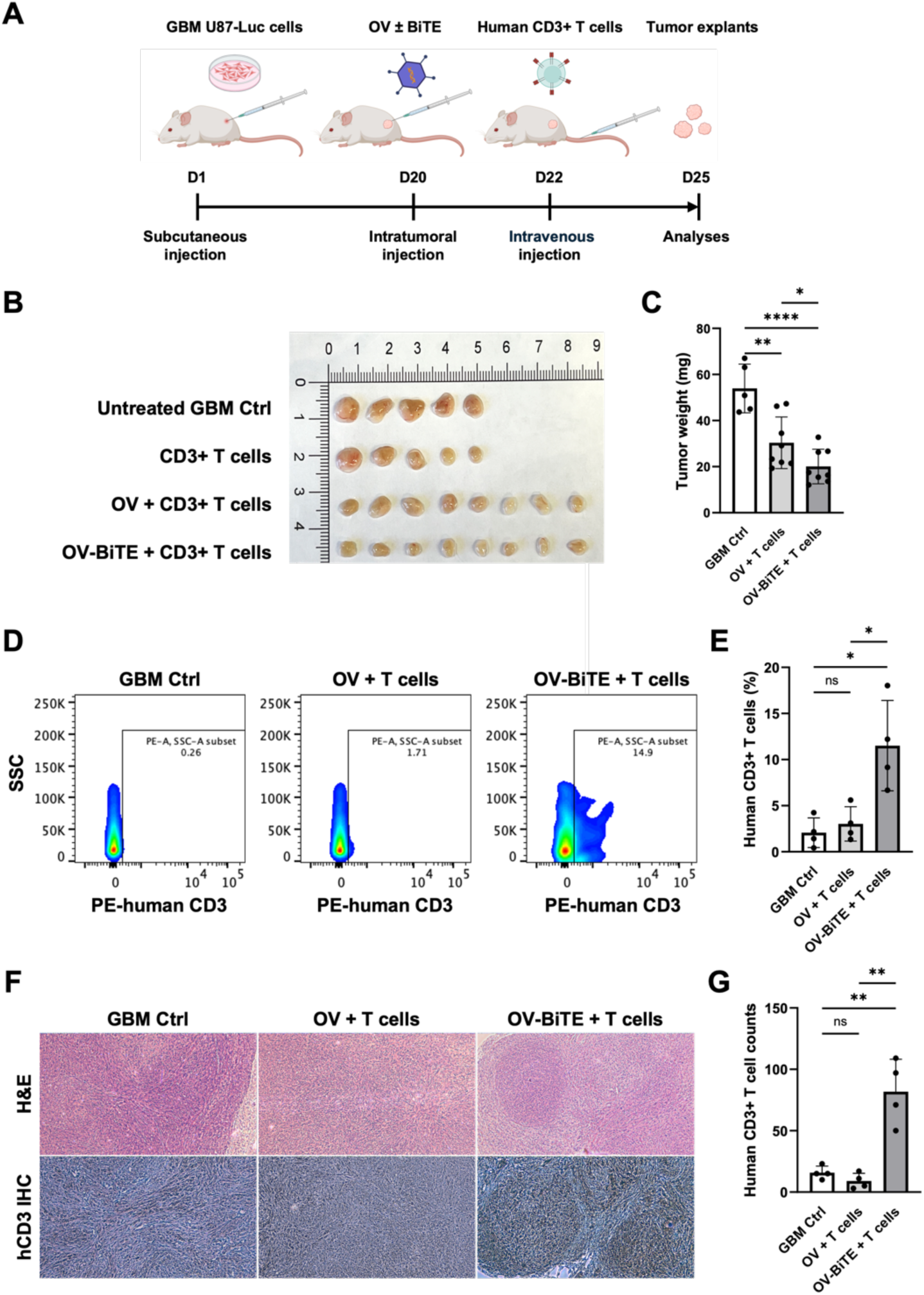
In vivo T cell recruitment assay of Adenoviral protein BiTE. GBM tumor mouse model established with U87-luc cells (A). OV were intratumorally injected and 2 days after adenovirus injection, CD3+ T cells were injected by tail vein and on day 25, all tumors isolated from mice and their size was analyzed (B and E). Human CD3+ T cell population in tumor tissues were analyzed by FACS (C and D). Also, hCD3 + cells were detected by IHC analysis (F). Each value represents the mean ± S.D. for three separate experiments. *p < 0.05, **p < 0.01 and *** p < 0.001

### Combination of OV-BiTE & CAR-T cells effectively eliminates GBM tumor in mice

Based on the abovementioned *in vitro* GBM killing efficacy of OV-BiTE and CAR-T cells, as well as the enhanced T cell recruitment by OV-BiTE, we proceeded with combination therapy in a mouse GBM tumor xenograft model (Figure 5A). All mice were subcutaneously injected with the GBM U87-Luc cells on the right flank. Twelve days later, adenovirus was intratumorally injected as described elsewhere ^52^. Another 3 days later, human CD8+ T cells or therapeutic CAR-T cells were infused through tail vein injection. On the last day of experiments, mice were euthanized, and tumor tissues were isolated. Moreover, GBM tumor growth in different treatment groups was monitored at indicated intervals from Day 0 to Day 31 by using a luminescence *In Vivo* Imaging System (IVIS) (Figure 5B). There were no body weight losses from the mice throughout the entire duration of the therapy (Figure 5C) and the luminescence intensity of the region of interest (ROI) was analyzed for each mouse (Figure 5D). On Day 24 and 31, all three OV-BiTE & CAR-T groups showed significant reductions in tumor luminescence intensity compared to CD8+ T cell treated group (Figure 5E and F), where OV-BiTE with combi CAR-T group exhibited even greater efficacy than OV-BiTE with either EGFR or EGFRvIII CAR-T alone (Figure 5G and H). These results were further validated by assessing the size and weight of explanted tumor masses (Figure 5I-L). To further establish the GBM killing effects of our multipronged immunotherapy, we performed immunohistochemistry (IHC) on all tumor explants. Hematoxylin and eosin (H&E) staining revealed that the cellularity, as indicated by hematoxylin purple blue staining of nuclei, in untreated GBM control and CD8+ T cell treated groups was visibly higher than that in the three groups received OV-BiTE & CAR-T cell treatment (Figure 6A). IHC staining of GBM specific tumor associated antigen IL13R⍺2 was significantly higher in the untreated GBM control and CD8+ T cell treated groups than in the immunotherapy groups (Figure 6B and C). IHC staining of human CD3+ T cells was significantly higher in the immunotherapy groups than in the untreated GBM control and CD8+ T cell treated groups (Figure 6B and D). In summary, we have demonstrated that sequential application of OV-BiTE and CAR-T cells can effectively eliminate GBM tumor in our mouse model, and that OV-BiTE followed by combi CAR-T cells exhibiting even greater efficacy than OV-BiTE with either EGFR or EGFRvIII CAR-T alone.

**Figure 5.**
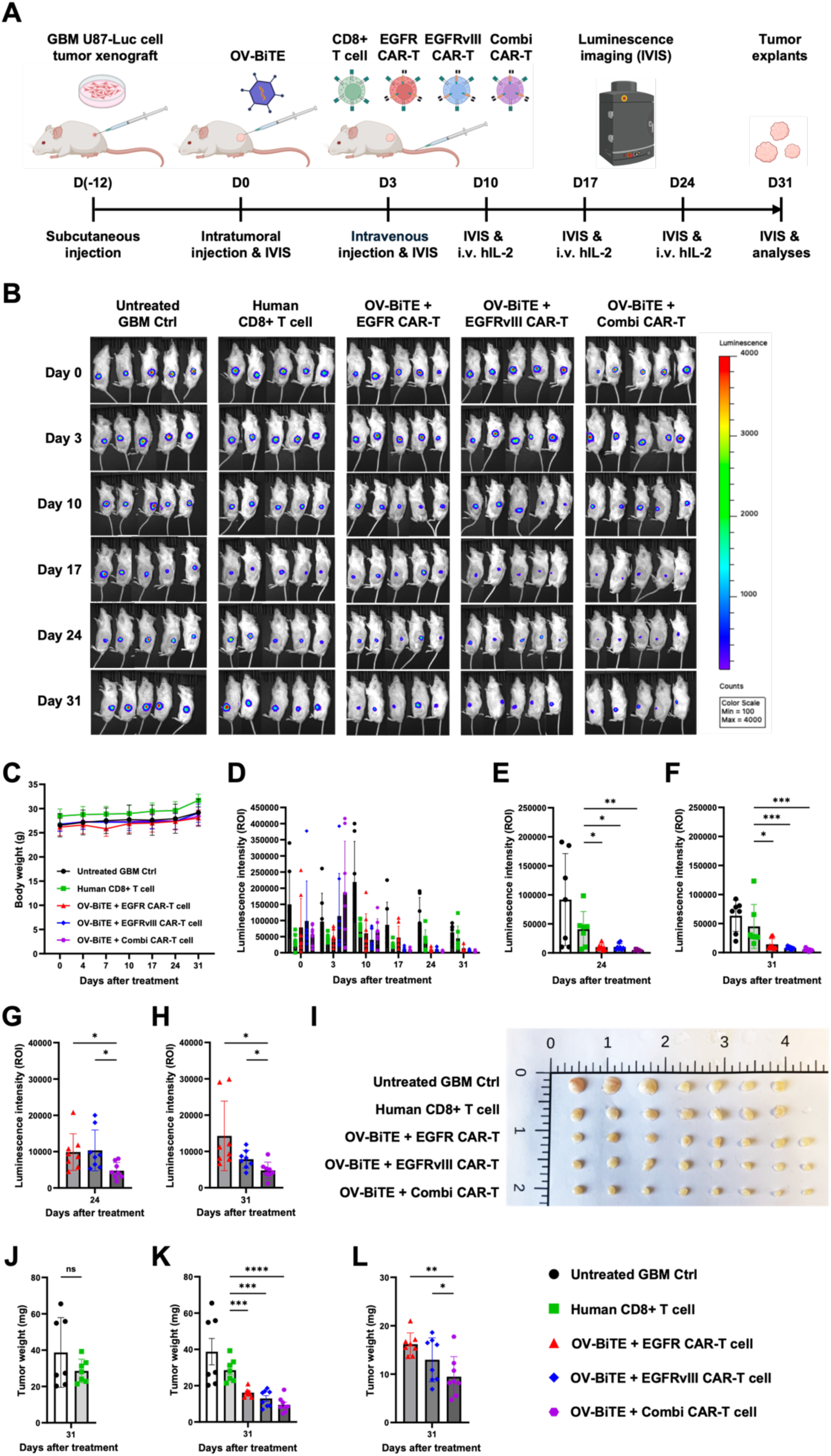
In vivo combination immunotherapy of CAR-T cells and OAV. All animal experimental processes were explained (A). During the experiment, tumor-bearing mice were analyzed using the IVIS imaging system (B). At the endpoint of the experiments, all mice were sacrificed, and tumor tissues were isolated from the mice (I). Luminescence intensity of tumors was analyzed using Live imaging software (E-H). Body weight was measured twice a week during the experiments (C), and tumor weight was compared between different experimental groups (J-L). Each value represents the mean ± S.D. for three separate experiments. *p < 0.05, **p < 0.01, ***p < 0.001, and ****p < 0.0001.

**Figure 6.**
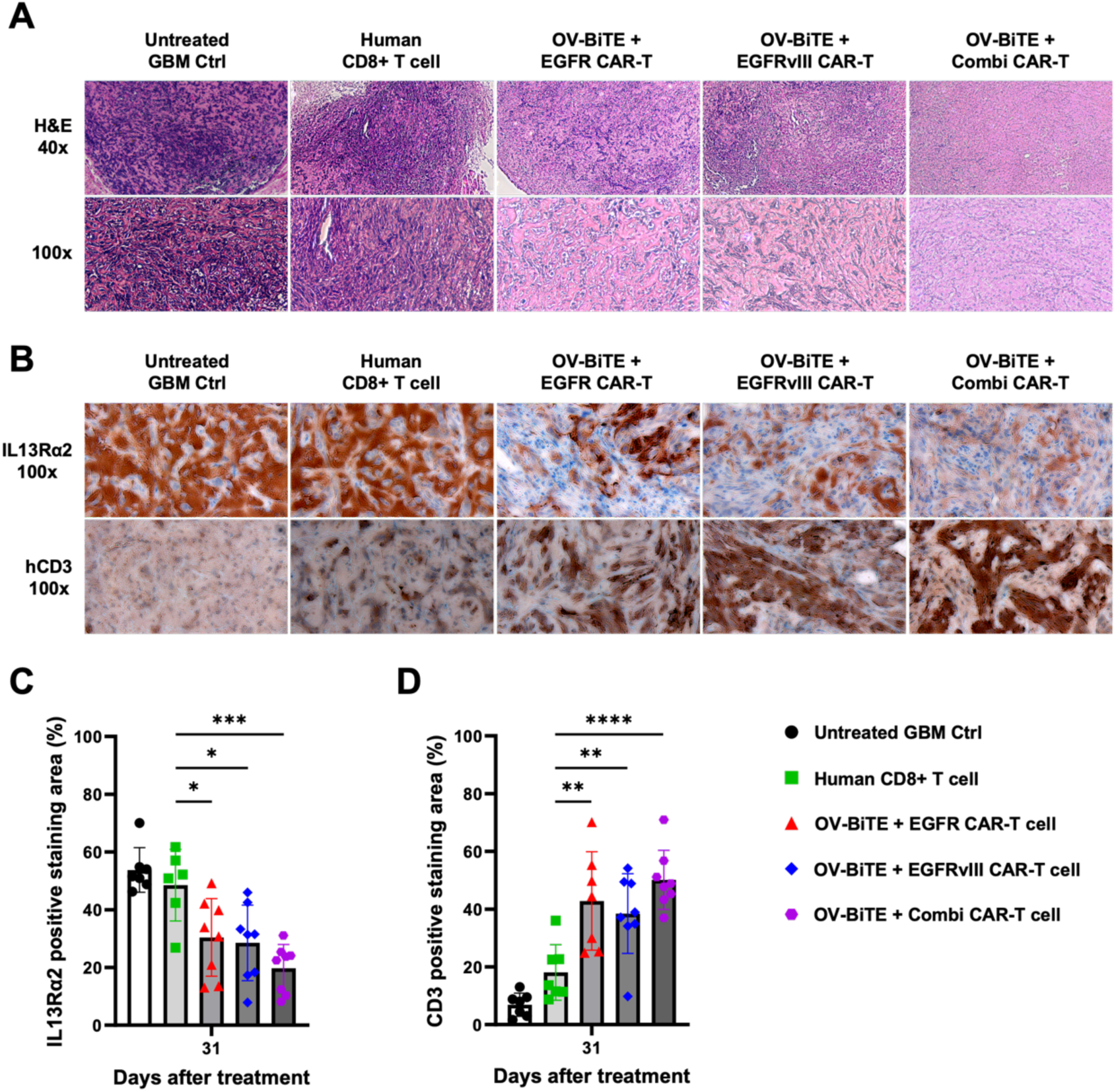
Immunohistochemistry analysis of tumor tissues. H&E staining sections were analyzed for tumor cell density in all groups (A). And IL13R⍺2 and hCD3 were detected in tumor tissues and their positive percent area was analyzed (C and D). Each value represents the mean ± S.D. for seven separated tumor tissues in control and hCD8+ Tc group and eight separated tumor tissues in three therapeutic groups. *p < 0.05, **p < 0.01, and ***p < 0.001

## Discussion

GBM is one of the most difficult challenges in oncology, with its immunosuppressive tumor microenvironment posing significant barriers to effective therapy. Conventional approaches such as radiotherapy and chemotherapy have demonstrated limited success in extending survival, underscoring the need for innovative approaches. Immunotherapy, particularly CAR-T cell therapy, has emerged as a promising modality. Recent clinical trials utilizing CAR-T cell therapies in adult and pediatric high-grade gliomas have demonstrated safety and feasibility of intracranial CAR-T cell delivery, and provided evidence of transient responses, supporting further investigations of this transformative approach to treating a formidably aggressive cancer ^22–24,53–55^. Currently, there has been no universal CAR-T cell target for GBM identified yet. Hence, combinations of targets and engineered CAR-T cells expressing immunostimulatory molecules to enhance their efficacy should continue to be developed. Our current study intends to fill such a gap by introducing an innovative multimodal approach toward overcoming the challenges associated with CAR-T therapy in solid tumors. By intratumorally administering an engineered oncolytic adenovirus to overexpress a BiTE, we could demonstrate a significantly enhanced recruitment of endogenous as well as therapeutic T cells to the tumor. This approach aimed to remodel the tumor microenvironment by converting immunologically ‘cold’ tumors, characterized by low immune activity, into ‘hot’ tumors with enhanced immune activation, thereby improving therapeutic efficacy. The BiTE was designed with two arms, one for IL13R⍺2 that is specifically overexpressed on GBM cells and the other for CD3 of T cells. Through BiTE-expressing OV, we expect that more cell lysis in tumor tissue and more immune activity from T cells can enhance the therapeutic effect of CAR-T cells.

It is well-established that both lytic and non-lytic cytokines are involved in the antitumor activities by CAR-T cells ^56^. As part of the *in vitro* cytotoxicity evaluation, our CAR-T cells were cocultured with two different GBM cell lines at appropriate effector-to-target (E:T) ratios, and these engineered CAR-T cells had significantly higher cytokine production than that induced by primary CD8+ cells. Notably, the secretion of lytic cytokines, e.g., perforin and granzyme B, which are involved in direct apoptosis of cells, was particularly elevated. These results were further validated through antitumor toxicity evaluations using GBM cells and GBM spheroids. Differences in tumor antigen expression between the cell lines may account for the variability in significance among the CAR-T groups. To evaluate whether CAR-T cells could overcome the physical limitations of the tumor microenvironment, we established a GBM–blood– brain barrier (BBB) spheroid model constructed by layering GBM core spheroids with astrocytes, endothelial cells, and pericytes. CAR-T cells induced significant tumor cell death even within the inner regions of the BBB spheroids, suggesting their ability to infiltrate and remain cytotoxic beyond the BBB-like barrier.

*In vivo* tumor xenograft model was used to evaluate the functionality of BiTE. Mice treated with OV-BiTE showed significantly inhibited tumor growth, as shown by reduced tumor size, vascularization, and weight. And enhanced CD3^+^ T cell infiltration was observed, likely due to BiTE screetion following adenovirus infection and lysis, facilitating cell migration into tumor. In the *in vivo* therapy experiments, both single CAR-T and combi CAR-T cells showed positive results, with the same number of CAR-T cells but different anticancer therapeutic efficacy. The two single CAR-T therapy groups have no differences in tumor growth inhibition, but the combi CAR-T cell group showed significant differences compared to the two single CAR-T cell groups. The observed results can be attributed to the highly heterogeneous nature of GBM ^57–59^, where a single tumor cell expresses multiple tumor antigens. This suggests that utilizing two or more CAR-T cells targeting different antigens is more effective than either one for GBM treatment. Furthermore, H&E histology and immunohistochemistry revealed a significant reduction in overall cellularity, decreased tumor antigen IL13R⍺2 expression but increased CD3+ cell staining in the tumor explants of OV-BiTE & Combi CAR-T group. In addition, we observed no side effects such as body weight loss or hair loss in animals during the therapy.

In summary, our study has demonstrated robust antitumor efficacy of the multipronged OV-BiTE & CAR-T modality in preclinical GBM models. The potential clinical applications of this multimodal platform may address key challenges associated with CAR-T cell immunotherapy in solid tumors, such as the immunosuppressive tumor microenvironment, thus, extending CAR-T cells’ utility beyond hematologic malignancies, where it has shown remarkable success. Moreover, the methodologies described in this study lay the groundwork for a novel combination therapeutic platform that synergizes the unique mechanisms of action of oncolytic viruses and cell-based immunotherapies. This approach can potentially transform the therapeutic landscape for solid tumors, providing a versatile framework for future translational studies and clinical applications. We anticipate that the principles and strategies presented here will contribute to the broader development of advanced oncolytic virus-based Immunotherapeutics for cancer treatment.

### Limitations of the study and Future directions

One of the primary limitations of our study is the use of a subcutaneous tumor model rather than an orthotopic model that more accurately mimics the brain microenvironment where GBM naturally arises. The subcutaneous model, while widely used for its experimental convenience, lacks the complex interactions present in the central nervous system (CNS), such as the influence of the BBB, CNS-resident immune cells, and the unique tumor microenvironment of GBM. Consequently, the findings of this study may not fully reflect the therapeutic dynamics and efficacy of OV-BiTE in an authentic GBM context. To address these limitations, future studies will focus on evaluating the antitumor activity of OV-BiTE in orthotopic models using humanized mice. This approach will enable a more accurate assessment of the therapy’s interactions with other immune cell populations and its efficacy within the CNS-specific tumor microenvironment. Such models will be instrumental in bridging the gap between preclinical findings and their translation to clinical applications, providing deeper insights into the therapeutic potential of OV-BiTE for GBM treatment. Furthermore, validation experiments were performed using U87 and U251 cell lines, and the use of these legacy cell lines share little resemblance to patient glioblastoma. The purpose of the current study is to provide proof of principle in combining OV-BiTE and CAR-T cells, GBM patient derived cells may be used in future studies. In addition, in comparison with the current CAR-T cell therapy, CAR-NK cell therapy can use allogeneic cells, CAR-dependent and -independent tumor killing mechanisms, has low risk of graft-versus-host disease, cytokine release syndrome or neurotoxicity. Hence, future studies may investigate immunotherapies that combine armed OV with CAR-NK cells to treat GBM. Furthermore, future studies could be investigating the direct use of purified recombinant BiTE proteins as an alternative to viral delivery. While current attempt based on virus-mediated production, BiTE protein delivery may enhance control overdosing and reduced immunogenicity, and easier apply to clinical translation.

## Resource Availability

Lead contact: Olin Liang, Ph.D.

Materials availability

Data and code availability

## Acknowledgments

This work was supported in part by the National Institutes of Health P30GM145500 to O.D.L., by the Brown University Legorreta Cancer Center Translational Research Award to O.D.L., by the Brown Physicians Inc. Academic Assessment Research Awards to O.D.L. and P.B., and by the George Funds for Hematology and Oncology Research to O.D.L..

## Author Contributions

Conception and design, M.J.C. and O.D.L.; development of methodology, M.J.C., E.Y.S., B.A., Y.E.L., A.G.R., P.B., A.M.R., C.C.C., S.E.L., E.T.W. and O.D.L.; acquisition of data, M.J.C., E.Y.S., B.A., Y.E.L., and O.D.L.; analysis and interpretation of data, M.J.C., E.Y.S., and O.D.L.; writing, review, and/or revision of manuscript, M.J.C., E.Y.S., B.A., Y.E.L., A.G.R., P.B., A.M.R., C.C.C., S.E.L., E.T.W. and O.D.L.; administrative, technical, or material supports, E.Y.S., B.A., Y.E.L., A.G.R., P.B., A.M.R., C.C.C., S.E.L., and E.T.W.; study supervision, O.D.L..

## Declaration of interests

All authors declare no competing interests.

## Experimental model and detailed method

### Cell culture

All GBM cell lines and HEK293FT were obtained from the American Type Culture Collection (ATCC, Manassas, VA). Human peripheral blood mononuclear cells (PBMCs) were obtained from Lonza (Lonza Group AG, Switzerland). All GBM cell lines were cultured in DMEM/F12 media (Thermo Fisher Scientific, Waltham, MA) supplemented with 10 % Fetal Bovine Serum (FBS, Thermo Fisher Scientific), 1 % Antibiotic Antimycotic Solution (100X, Thermo Fisher Scientific). T cells were cultured in CTS^TM^ OpTmizer T-Cell expansion SFM (Thermo Fisher Scientific) supplemented with 10 % FBS, 1 % Antibiotic Antimycotic Solution, and L-glutamine (200 mM, Thermo Fisher Scientific). HEK293FT was used and cultured for the lentivirus packaging in DMEM media (Thermo Fisher Scientific) supplemented with 10 % FBS and 1 % Antibiotic Antimycotic Solution (Thermo Fisher Scientific). All lentivirus preparation and transduction for making the luciferase overexpressing cell lines was proceeded by Brown University Legorreta Cancer Center Cell & Genome Engineering Shared Resource.

### T cell expansion and Activation

CD8+ T cells were isolated from human PBMC using EasySep Human CD8+ T Cell Isolation Kit. After isolation, all T cells were counted with TC20 automated cell colony counter (Bio-Rad, Hercules, CA). For the expansion, Dynabeads^TM^ Human T-Activator CD3/CD28 for T cell expansion and activation was added to the CD8+ T cells and cultured with IL-2 (PEPROTech, 100IU/mL; Cranbury, NJ) in complete CTS^TM^ OpTmizer T-Cell expansion SFM (Thermo Fisher Scientific). To activate the T cell before transduction of Lentivirus, increase the amount of Dynabeads to double the T cell number (1:1 for expansion and 2:1 for activation).

### Generation of CAR-T cells

The 293FT cells (Thermo Fisher Scientific) were used for the lentivirus packaging. All vector plasmids encoded CAR were designed in our lab, and CAR plasmid was produced by Creative Biolabs (NY, US). For lentivirus production, 2 million 293FT cells were seeded to a 10 cm culture dish and incubated 24 hr in 10 % DMEM (Thermo Fisher Scientific). Moreover, 1 day later, four types of packaging vectors (gag/pol, Tet, Rev, and VSVG) and expression vectors were mixed with LipoD293 *In Vitro* DNA Transfection reagent and incubated the mixture for 15 minutes at room temperature (RT). The culture media included mixture was changed with 10 mL of fresh media after 24 hours of transfection. At 48 hr and 72 hr after transfection, Lentivirus, including culture media, was gathered and stored at 4 degrees. Lentivirus soup was centrifuged to remove cells and debris, and the supernatant was transferred to a new tube and filtered with 0.45 µm syringe filters. For the concentration of Lentivirus, add Lenti-X Concentrator (Takara, Japan) at a 1:3 ratio and incubate for 2 hr on ice. After incubation, the virus concentrate mixture was centrifuged at 1500 g for 1 hr 30 min. After centrifugation, remove the supernatant and suspended lentivirus pellet with HBSS solution (Thermo Fisher Scientific).

Add Lentivirus at a MOI of 10 to a new microtube for the lentivirus transduction to T cells. Add polybrene at eight µg/mL of concentration and incubate for 15 min. After incubation, a lentivirus mixture was added to the prepared CD8+ T cells. For the infection, centrifuge lentivirus-treated T cells at 700 g, 45 min, 32 °C. The cells were centrifuged at 1200 rpm one day after transduction, 3 min. Moreover, add fresh media including 100 IU/mL of IL-2.

### Elisa and EliSPOT for the cytokine assay

Seed the 2*10^5 two GBM cell lines (U251 and U87) in 6 well culture plates with 2 mL culture media. One day after seeding, remove supernatant and add 1 mL complete culture media of GBM cell lines and add activated T cells (CD8+ T cells and three types of CAR-T cells) in 1 mL T cell culture media (1:1 ratio) and incubate for 24 hr in 37-degree cell culture incubator. Two GBM cell lines, CAR-T cells and T cells (human cd8+ T cell), and their cytokine release profiling were analyzed by ELISA and Elispot.

### Western Blot

GBM cell lines and astrocytes were lysed using RIPA buffer (Thermo Fisher Scientific) supplemented with a protein inhibitor. Cell lysates were incubated on ice for 10 min, and protein samples were collected by centrifugation at 14,000 rpm for 10 min. Total protein was quantified with BCA assay. All samples were separated using SDS-PAGE and transferred onto a nitrocellulose membrane using a Trans-Blot Turbo transfer system (Bio-Rad). Transferred membranes were blocked with 3 % skim milk in TBST buffer for 2 hr, RT. After blocking, add a primary antibody in fresh skim milk overnight at 4 °C. A fluorescence-labeled secondary antibody was added to the washed membrane for the detection. The protein on the membrane was detected using Odyssey (LI-COR, Lincoln, NE). The following antibodies were used for tumor antigen detection: anti-beta actin (4970S), anti-IL13RA2 (85677S), and anti-EGF Receptor (54359S) from CST (Danvers, MA). Anti-EGFRviii (XJ3728096) from Invitrogen (Waltham, MA) and anti-6X His tag (EPR20547) from Abcam (Cambridge, UK).

### Flow Cytometry

CAR-T cells were stained using DAPI solution (BioLegend, San Diego, CA), PE anti-human CD3 (OKT3), APC anti-human CD4 (RPA-T4), Alexa Fluor 700 anti-human CD8 (SK1), and anti-human IgG Fc (M1310G05) were obtained from BioLegend. DAPI solution, DPBS (Gibco)

For the cell staining, the cells for analysis were washed with DPBS three times, 500 µL PBS, including the target antibody, and incubated at 4 °C for 1 hr. After incubation, the cells were washed three times with DPBS and added 300 µL of fresh DPBS, including the DAPI solution. Cell acquisition and analysis were performed using an LSR II (BD, Franklin Lakes, NJ). For the data analysis, FlowJo software was used.

### *In vivo* mouse xenograft model

Human glioblastoma U87-MG-luciferase cells were cultured in DMEM/F12 supplemented with 10 % fetal bovine serum and 1% Antibiotic-Antimycotic. Cells were harvested at 80-90 % confluency, washed with PBS two times, and resuspended in a 1:1 mixture of serum-free DMEM and Matrigel matrix (Corning) at a concentration of 5 × 10⁶ cells/200μL. Six-week-old male NSG(NOD/SCID) mice were anesthetized with isoflurane, and 200 µL of the cell suspension was injected subcutaneously into the right flank. Tumor growth was monitored two times a week using calipers, and tumor volume was calculated using the formula: (length × width²)/2. When tumors reached approximately 100 mm³, OV-BiTE and CAR-T cell treatment commenced, as detailed below.

### OV-BiTE production

To prepare oncolytic adenovirus, all procedures followed the manufacturer’s instructions (The AdenoQuick2.0 Cloning System Manual, OD260 Inc.). First, digest the pAd1127, pAd1128, pAd1129, and pAd1130 plasmid derivatives with *Sfil* ligate the digested plasmid with each other and package the ligation product into phage ƛ, and infect competent cell with the packaged phage ƛ. To prepare cosmid DNA, digest 15 μg cosmid DNA with *PacI*. Verify the completion of the digestion and the quality of the DNA by agarose gel electrophoresis. Precipitate the DNA with EtOH, centrifuge, and resuspend the DNA pellet in 60μL sterile TE pH 7.5. Then, adenoviral cosmid was transfected into AdenoX 293FT packaging cells, followed by instructions. For the identification and titration of adenovirus, the Adeno-X GoStix and Adeno-X rapid titer kit (Takara) were used. All procedures followed their instructions. For the *in vitro* test, all OVs were treated to cancer cells at a MOI 100. All adenovirus experiments were conducted in accordance with Institutional Biosafety Committee of Lifepan (Protocol Number: #502223)

### OV-BiTE and CAR-T cell injection

*In vivo*, combination therapy, on day 0, MOI 20 OV-BiTE (100 million virus particles) was intratumorally injected in 100 μL PBS. Mice baring subcutaneous glioblastoma tumors were anesthetized with isoflurane, and the tumor area was sterilized with 70% ethanol. Using a 30-gauge needle, 100 µL of the OV-BiTE solution was injected intratumorally at three separate sites to ensure even distribution within the tumor mass. Three days after OV-BiTE injection, 5 million of prepared CD8+ T cells and CAR-T cells (E: T raion = 1:1) were intravenously injected by xenograft mice tail vein under light isoflurane anesthesia. Post-injection, animals were observed for 1 hour to ensure recovery from anesthesia and evaluated for therapeutic response at predetermined time points. And tumor volume was measured twice a week. All animal experiments were conducted in accordance with institutional guidelines and approved by the Institutional Animal Care and Use Committee (IACUC) of Rhode Island Hospital (Protocol Number: #504123). Efforts were made to minimize animal suffering, including the use of appropriate anesthetics and humane endpoints. The facility operates in compliance with the Guide for the Care and Use of Laboratory Animals published by the National Research Council.

### Immunohistochemistry assay

Formalin-fixed paraffin-embedded (FFPE) tissue sections (4 µm) were mounted on positively charged slides and air-dried overnight. Slides were baked at 50-60 °C for 1 hour, deparaffinized through graded xylene and ethanol series, and subjected to antigen retrieval using EnVision FLEX Target Retrieval Solution, High pH (Agilent, K800021-2) in a pressure cooker (BioGenex, NW001-PC) at full microwave power for 10 min. Endogenous peroxidase activity was quenched using Peroxidase-Blocking Reagent (Agilent, K800021-2) for 10 min at room temperature. Non-specific binding was blocked with 5% non-fat milk for 30–60 min, followed by incubation with primary antibodies diluted in DAKO Antibody Diluent (Agilent, S0809) overnight at 4 °C. The following primary antibodies were used: anti-human CD3 (Abcam, ab17143; 1:200), anti-human CD8 (Abcam, ab245118; 1:250), anti-p53 (Agilent, IR61661-2; RTU), anti-alpha-SMA (Agilent, M085101-2; RTU), anti-IL13Rα2 (Abcam, ab250044; 1:100), and anti-His tag (Abcam, ab213204; 1:200). After washing, sections were incubated with EnVision HRP-conjugated secondary antibody (Agilent, K4061) for 45 min at room temperature. Visualization was achieved with DAB+ substrate-chromogen (Agilent, K800021-2) for up to 10 min. Slides were counterstained with Gill’s hematoxylin, blued with ammonia water, dehydrated, and mounted with coverslips.

## Statistical analysis

All statistical analyses were performed using GraphPad Prism version 10 (GraphPad Software, San Diego, CA). Data are presented as mean ± standard deviation (SD). Comparisons between the two groups were conducted using unpaired t-tests. For comparisons among multiple groups, one-way ANOVA followed by Tukey’s post hoc test was employed. Survival curves were generated using the Kaplan-Meier method and compared using the log-rank (Mantel-Cox) test. A p-value of less than 0.05 was considered statistically significant.

## Supplementary Figures

**Supplement Figure 1.**
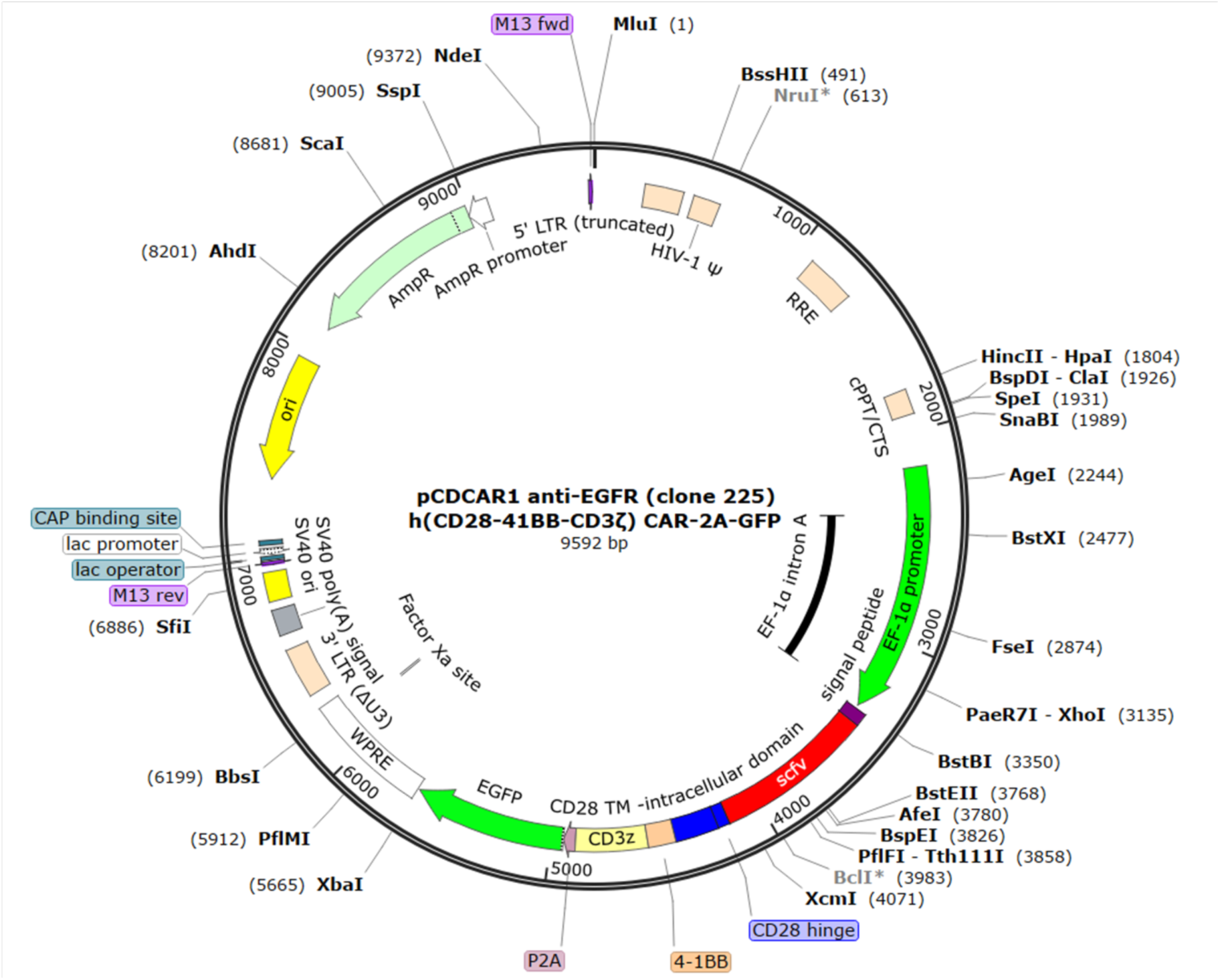
Genetic mapping of anti-EGFR CAR expression plasmid DNA. All CAR constructs have same parts except ScFv part for each CAR target. And it showed general third generation CAR constructs including CD3z, CD28, and 4-1BB. And all pCAR were designed with EGFP.

**Supplement Figure 2.**
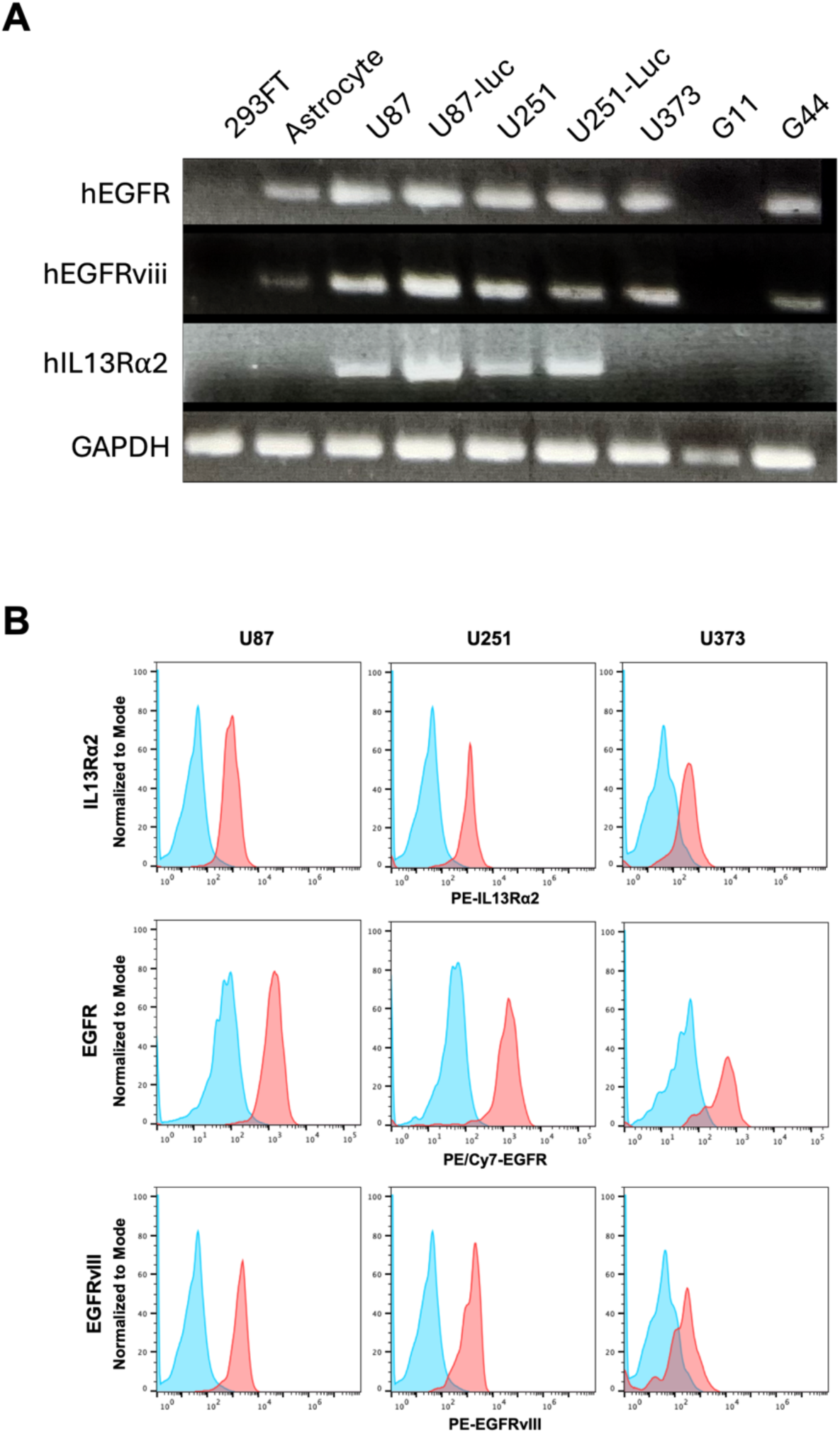
Tumor antigen expression of various GBM cells. The RNA of human kidney cell 293FT cell, human Astrocyte, U87, U87-luc, U251, U251-luc, U373, G11 (GBM patient derived cells), and G44 (GBM patient derived cells) were prepared and four targets expression including housekeeping gene GAPDH was analyzed qPCR (A). And surface antigen expression (IL13R⍺2, EGFR, and EGFRviii) of three GBM cell line was detected by FACS (B).

**Supplement Figure 3.**
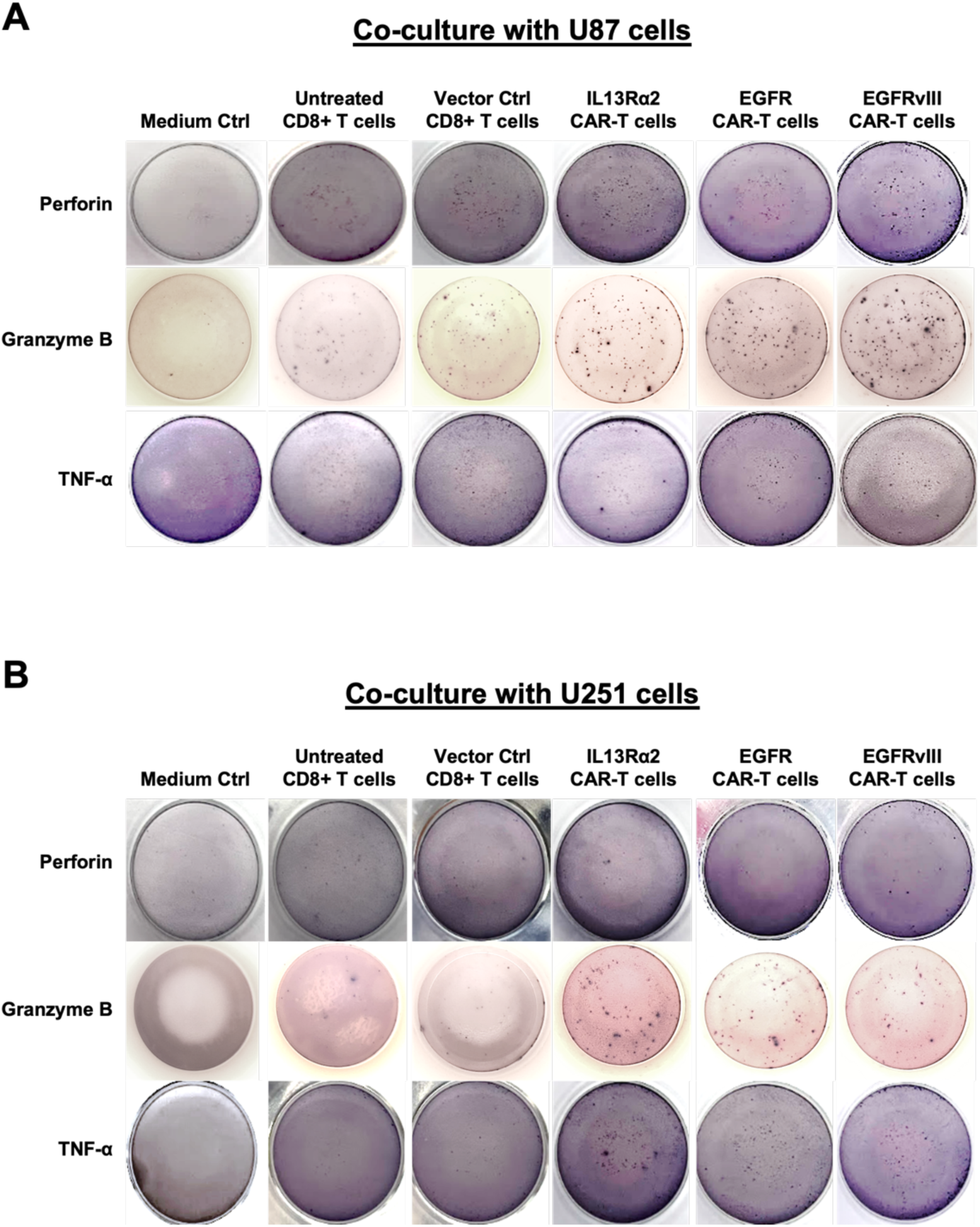
ELISPOT analysis results images about human Granzyme B and Perforin. Three different CAR-T cells were treated to U87 (A) and U251 (B) and after incubation the positive purple colored crystal in the each ELISPOT plate well was detected.

**Supplement Figure 4.**
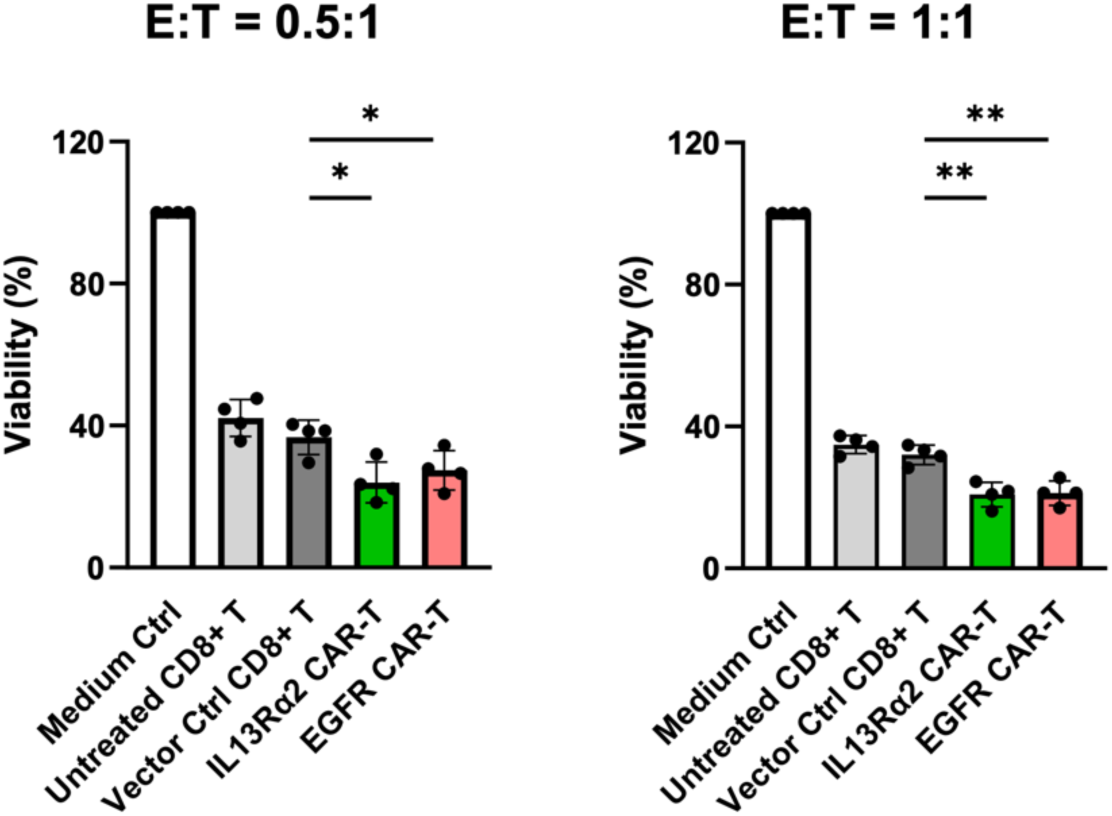
Identification of E: T ratio for killing assay of CAR-T cells. U87-luc cells were treated with control CD8+ T cell and two different CAR-T cells (EGFR and IL13R α2) at two E: T ration (0.5: 1 and 1: 1). And the cell viability was analyzed by luciferase activity. Each value represents the mean ± S.D. for three separate experiments. *p < 0.05, ** p < 0.01, and *** p < 0.001

**Supplement Figure 5.**
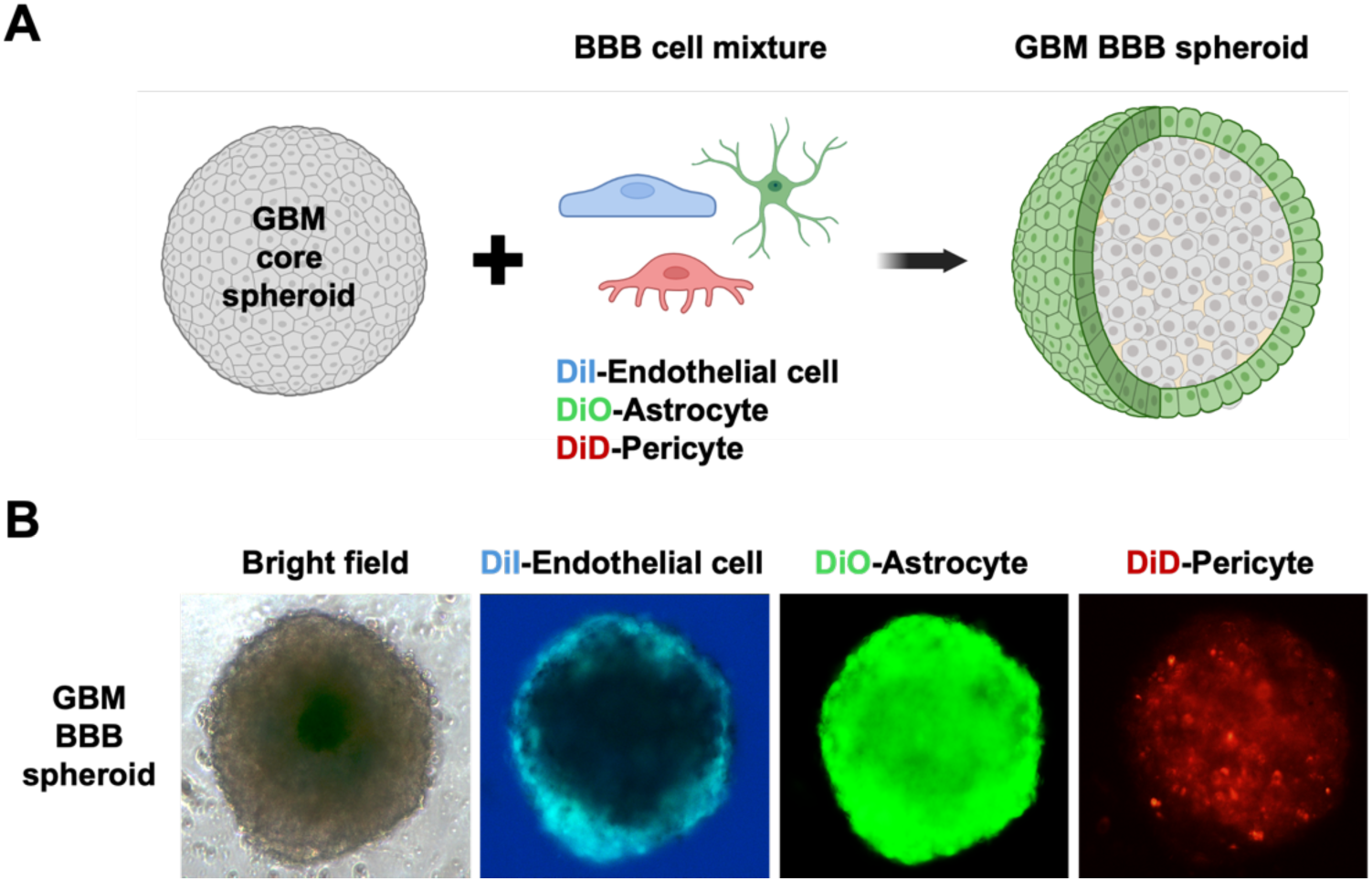
Development of GBM-BBB Spheroids. For evaluation T cell infiltration and killing in GBM-BBB Spheroids, the GBM core spheroid mixed with different fluorescent labeled three different types of BBB cells (A). The GBM-BBB spheroid formed by hanging drop method and location of each cell was detected with fluorescence microscope (B).

**Supplement Figure 6.**
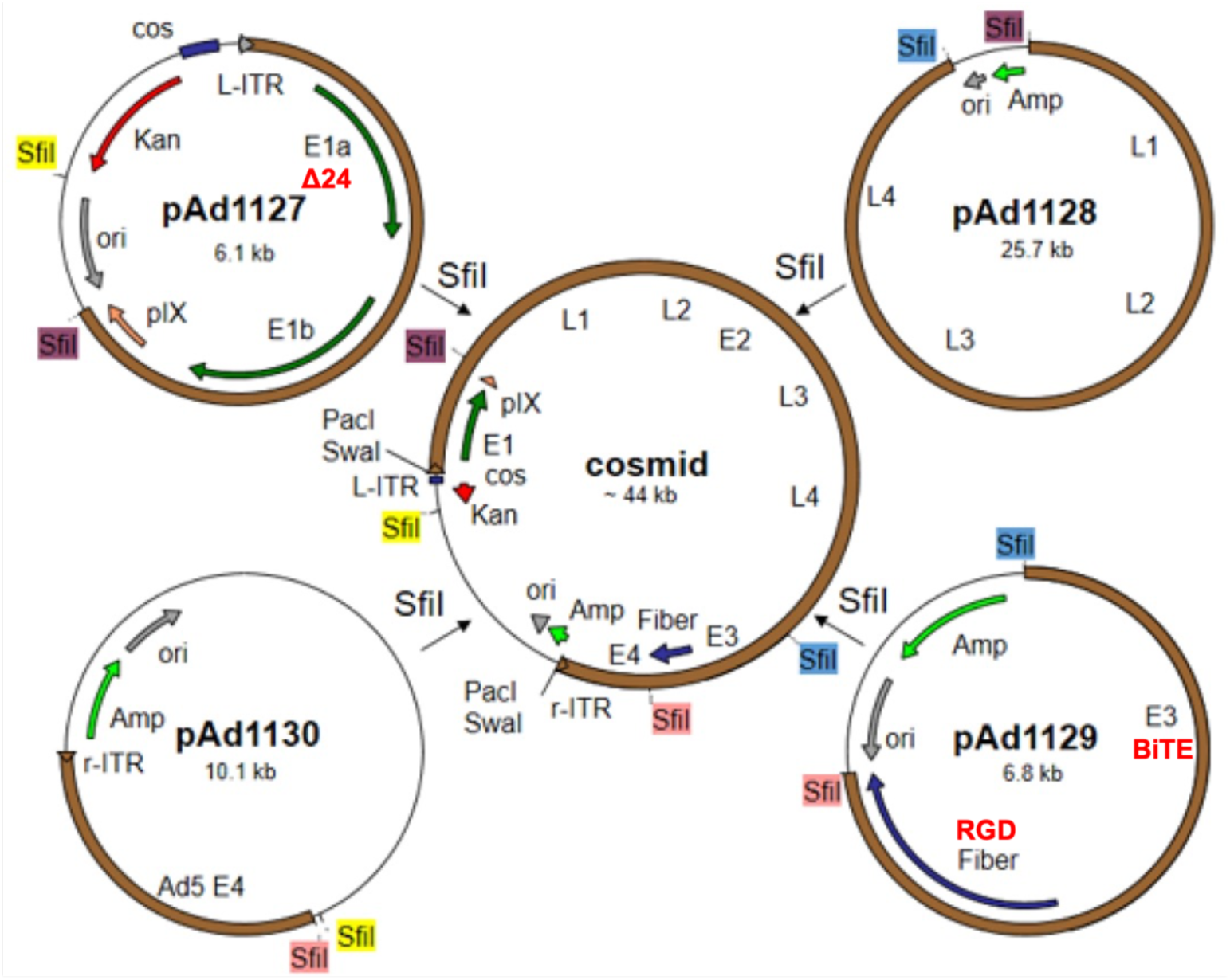
Adenoviral sub-plasmids and cosmid construct. All adenoviruses were produced by adenoviral cosmids. For making the adenoviral cosmid, four different types of plasmids combined. All adenoviruses have delta 24 modified E1A from pAd1127 and BiTE and RGD parts from pAd1129.

## Notes

### Competing Interest Statement

The authors have declared no competing interest.

